# Subdivision of ancestral scale genetic program underlies origin of feathers and avian scutate scales

**DOI:** 10.1101/377358

**Authors:** Jacob M. Musser, Günter P. Wagner, Cong Liang, Frank A. Stabile, Alison Cloutier, Allan J. Baker, Richard O. Prum

## Abstract

Birds and other reptiles possess a diversity of feather and scale-like skin appendages. Feathers are commonly assumed to have originated from ancestral scales in theropod dinosaurs. However, most birds also have scaled feet, indicating birds evolved the capacity to grow both ancestral and derived morphologies. This suggests a more complex evolutionary history than a simple linear transition between feathers and scales. We set out to investigate the evolution of feathers via the comparison of transcriptomes assembled from diverse skin appendages in chicken, emu, and alligator. Our data reveal that feathers and the overlapping ‘scutate’ scales of birds share more similar gene expression to each other, and to two types of alligator scales, than they do to the tuberculate ‘reticulate’ scales on bird footpads. Accordingly, we propose a history of skin appendage diversification, in which feathers and bird scutate scales arose from ancestral archosaur body scales, whereas reticulate scales arose earlier in tetrapod evolution. We also show that many “feather-specific genes” are also expressed in alligator scales. In-situ hybridization results in feather buds suggest that these genes represent ancestral scale genes that acquired novel roles in feather morphogenesis and were repressed in bird scales. Our findings suggest that the differential reuse, in feathers, and suppression, in bird scales, of genes ancestrally expressed in archosaur scales has been a key factor in the origin of feathers – and may represent an important mechanism for the origin of evolutionary novelties.

## Introduction

Feathers are a complex morphological structure that evolved in theropod dinosaurs (1, 2) (Fig. 1A), replacing scales across most of the body. However, extant birds retain scales on their legs and feet, which come in two principal types: overlapping “scutate” scales, present across most of the shank and foot (3), and tuburculate “reticulate” scales, found on the ventral foot and toe pads (4). The homology of feathers and scales has long been proposed based on similarities in development (5-9), which are greatest at the placode stage, when signaling between dermis and epidermis initiates feather or scale growth. In particular, feathers, bird scutate scales, and alligator scales all exhibit a thickened epidermal placode (9) and the expression of several placode-specific markers (7-9). Experiments with retinoic acid have also shown that ectopic feathers can be induced to grow on top of foot scales (10). However, feather and scale development differs dramatically after the placode stage. Whereas scale morphogenesis is relatively simple, feathers develop their branched structure via a unique and complex process of epidermal pattern formation (5, 6, 11) that produces the distinct morphology of the feather, including branching barbs, follicle, and deciduous sheath.

**Fig. 1.**
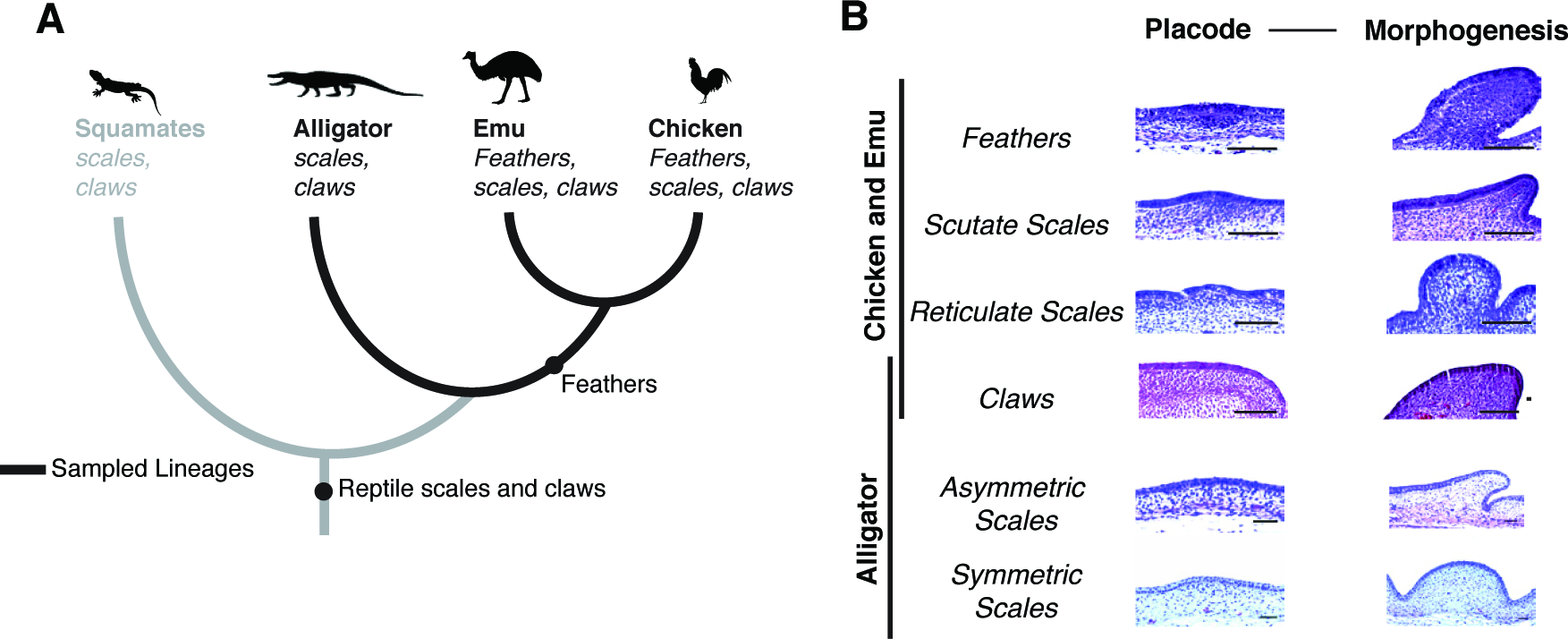
Phylogenetic history and sampling of reptile skin appendages. **(A)** Feathers evolved in the theropod dinosaur ancestors of birds, resulting in an addition to the skin appendage repertoire of birds. **(B)** Hematoxylin and eosin stained sections of skin appendage developmental stages sampled for RNAseq. Samples were collected at two stages of development, the placode stage when development is initiated, and later during their unique morphogenesis.

A key step in the evolution of feathers was the origination of a distinct regulatory program to orchestrate feather development. Over the previous two decades, investigations of feather development have yielded important insight into the signaling pathways and transcriptional regulators underlying feather development (11-18). However, how this regulatory program originated, and whether it is derived from that of another skin appendage, such as scales, is still unclear. Ideally, a full account of feather evolution should explain how the feather regulatory program is distinct from those of other skin appendages, how it originated from ancestral programs, and how birds have evolved the capacity to grow both feathers and scales from the skin of the same individual. This requires investigating not only the genetic program underlying feather development, but also that of other related skin appendages, in species that span the evolutionary events between ancestral scales and the evolutionary origination of feathers.

In this study, we investigate the evolution of the feather genetic program using mRNAseq to assay gene expression in different skin appendages of three archosaurs: chicken, emu, and alligator. The common ancestor of chicken (*Gallus gallus*) and emu (*Dromaeius novaehollandiae*) is the most recent common ancestor of all extant birds (Neornithes) (Fig. 1B), and thus represents the most distant possible comparison within birds. In chicken and emu, we sampled two embryonic feather tracts (dorsal and femoral), scutate and reticulate scales, and claws. In the outgroup species, the American alligator (*Alligator mississipiensis*), we obtained samples from asymmetric and symmetric body scales, as well as claws. Claws were included as they are a keratinized skin appendage with clear homology across reptiles, and because they allowed us to distinguish similarities between feathers and scales from those shared with all skin appendages. For each skin appendage, we sampled epidermis at the initiation of development – the placode stage – and at a second later stage during morphogenesis (Fig. 1B and Supplementary Table 1). We supplemented this with dermis collected for chicken skin appendages at both developmental stages.

Using both hierarchical clustering and phylogenetic analyses, we find that feathers and avian scutate scales likely originated from ancestral archosaur scales. Feathers and scutate scales continue to share many transcriptional and morphological similarities in birds that are not shared with reticulate scales or claws, which likely originated earlier in evolution. We document hundreds of new genes that are highly-specific to feathers, and show that many of them are expressed in distinct feather innovations. Notably, many genes with feather-specific expression patterns in birds are also expressed in alligator scales, plausibly representing ancestrally expressed genes that were recruited for new roles in feather development. This recruitment was accompanied by downregulation of feather-specific genes in bird scales. Based on this, we discuss a model of evolutionary diversification in which an ancestral structure gives rise to two descendant morphological structures through the reutilization and differential suppression of ancestrally-expressed genes.

## Results

### Feathers are more closely related to bird scutate scales and alligator scales than to other skin appendages

We investigated relationships among archosaur skin appendages using hierarchical clustering, parsimony evolutionary tree reconstruction, and t-distributed stochastic neighbor embedding (t-SNE) on developmental transcriptome data (19). For hierarchical clustering and t-SNE we analyzed a combined dataset containing gene expression from all three species. As a general outcome of these analyses, feathers, avian scutate scales, and alligator scales showed closer relationships to each other than to bird reticulate scales or claws (fig. 2 and fig. S1). In particular, hierarchical clustering at the two developmental stages always placed feathers *within* scales, such that they were more closely related to some scale types than others (figs. 2A,B). At the placode stage, we found feathers were most closely related to bird scutate and alligator scale placodes. A similar result was obtained from clustering morphogenesis stage skin appendages, except chicken scutate scales now clustered with chicken reticulate scales. Still, even during morphogenesis feathers maintained a close relationship with alligator scales and emu scutate scales. These clustering results were completely congruent with the parsimony phylogenetic reconstructions (fig. S1), which we conducted for each bird species separately for methodological reasons (19). Here, the close relationship between chicken feather and scutate scale placodes was independently supported by different gene sets in epidermis and dermis phylogenetic trees (fig. S2), indicating this relationship is not dependent on one, tissue-specific regulatory network, but represents a general pattern shared across different skin tissues.

**Fig. 2.**
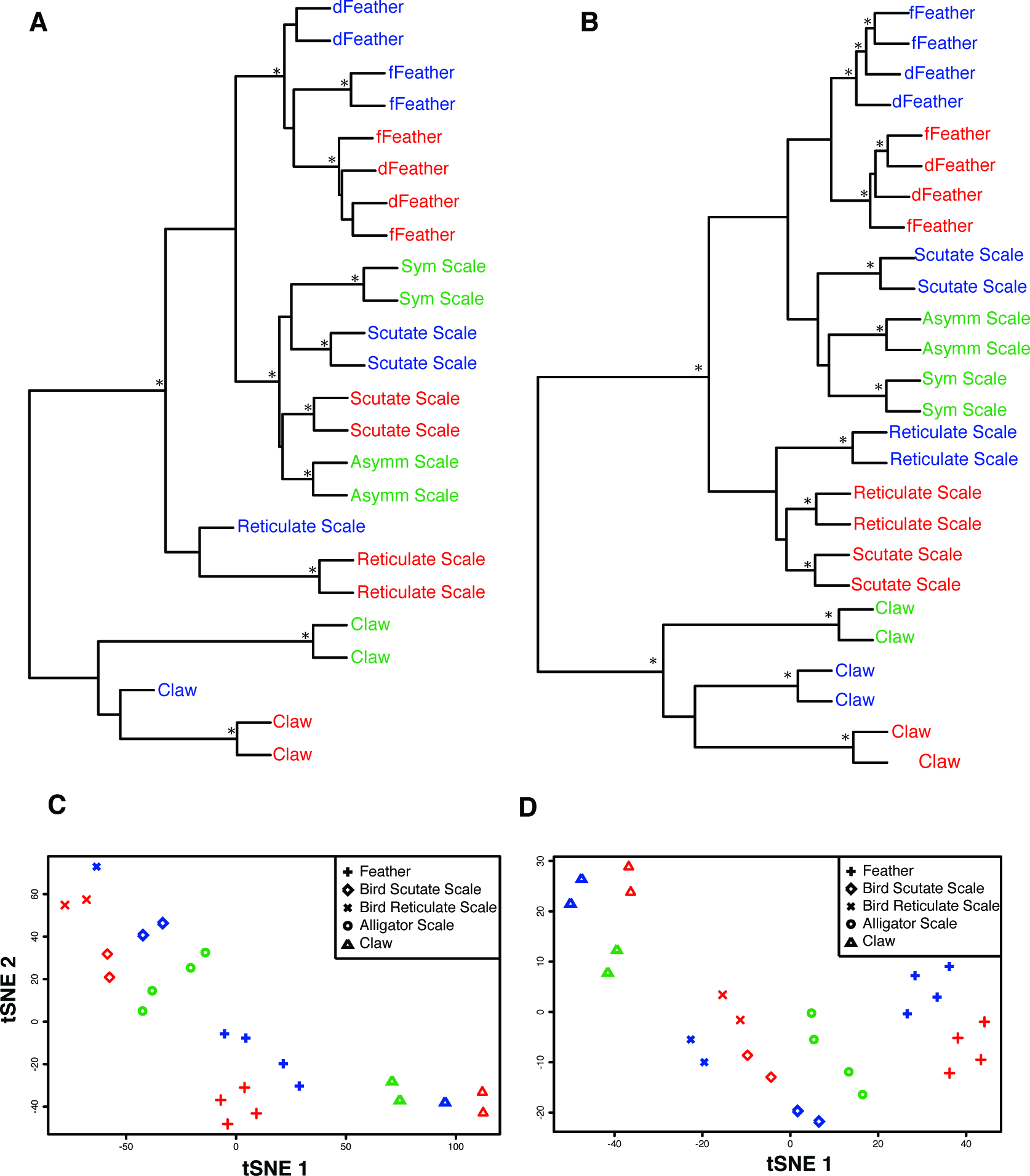
Clustering of skin appendage transcriptomes. **(A and B)** Hierarchical clustering of **(A)** placode stage and **(B)** morphogenesis stage skin appendage transcriptomes. **(C and D)** t-SNE plots of **(C)** placode and **(D)** morphogenesis stage transcriptomes. Chicken=red, emu=blue, and alligator=green, dFeather=dorsal feather, fFeather=femoral (chicken) or flank (emu) feathers, * > 80 bootstrap support.

As an alternative approach to determine tissue relationships we also used t-Distributed Stochastic Neighbor Embedding (t-SNE), a method that reduces high dimensional datasets to two dimensions, while preserving local relationships among samples. This revealed feathers and different scale types occur in a rough linear sequence across two-dimensional t-SNE space (figs. 2C,D), with feathers and reticulate scales at opposing ends of this spectrum. Alligator scale types were most similar to feathers at both stages of development, and were placed intermediate between scutate scales and feathers.

### Quantification of transcriptome covariance supports a close relationship between feathers and scutate scales

The evolution of a new morphological structure depends upon the ability to regulate gene expression independently of other closely related structures. This regulatory independence enables the new structure to maintain organ-specific gene expression, and to evolve organ-specific expression changes. We recently developed a method for quantifying correlated changes in gene expression between two tissues, which serves as a measure of their regulatory independence (20). Using a stochastic model of quantitative trait evolution, we estimate the level of correlated evolution (*γ*) between the gene expression programs of two tissues. The value of *γ* generally ranges between 0 and 1, with a value of 0 indicating expression changes are uncorrelated between the tissues, and a value of 1 indicating two tissues are completely correlated, and may share identical underlying gene regulatory networks.

Previously, we showed that placodes of feathers and scutate scales exhibit highly correlated evolution (*γ*=0.88, 95% CI [0.87, 0.89]; fig. 3 and table S1) relative to other bird and mammal tissues (20). This is consistent with our observations, and those of previous studies, that feathers and scutate scale placodes implement similar gene expression programs. Here, we extended this analysis to the morphogenesis stage, and found that the feather and scutate scales transcriptomes exhibit somewhat less correlated evolution during morphogenesis than they do at the placode stage (*γ*=0.84, 95% CI [0.83, 0.85]; fig. 3 and table S1). Still, even during morphogenesis, feather transcriptomes remain more highly correlated in evolution with scutate scale transcriptomes than with that of other avian skin appendages. In contrast, estimates of correlated evolution between other skin appendages did not change significantly during development. These observations suggest that feathers exert greater independent regulatory control in later development, yet likely continue to share common regulatory elements with maturing avian scutate scales.

**Fig. 3.**
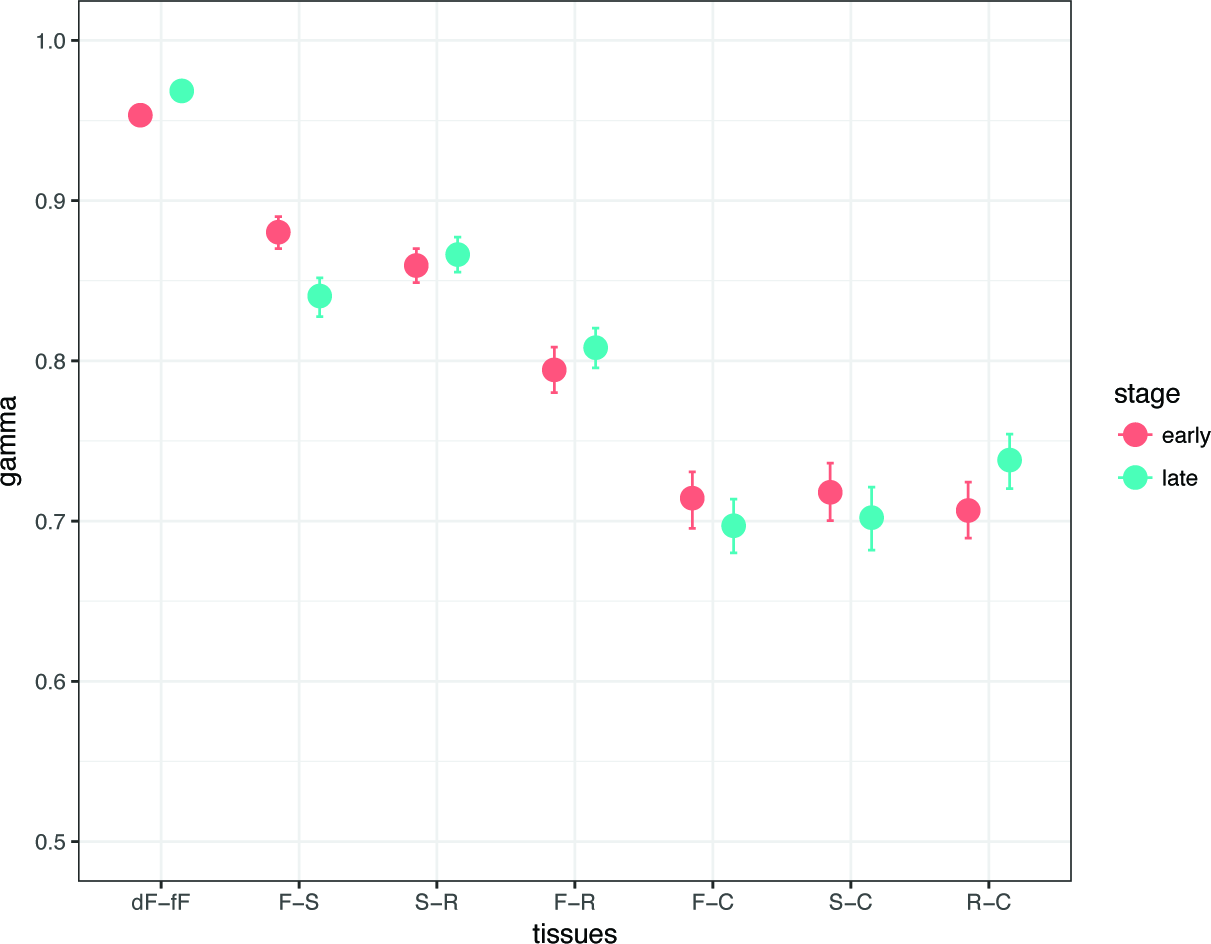
Evolutionary covariance of gene expression between different pairs of bird skin appendages. Covariance (γ) with 95% confidence intervals is an estimate of the average correlation of expression evolution across all genes between two tissues. C=claw, dF=dorsal feathers, fF=femoral feathers, F=feather samples grouped, R=reticulate scales, S=scutate scales.

### Identification of feather-specific genes

Our broad sampling of different avian skin appendages enabled us to identify genes that were highly-specific to feathers compared to other avian skin appendages. We identified 783 epidermal and 346 dermal genes with at least two-fold higher expression at the morphogenesis stage in feathers compared to all other avian skin appendages (fig. 4A). Feather-specific genes were enriched for transcription factors and signaling receptors (fig. 4B and table S2), confirming the importance of gene expression regulation in establishing differences among avian skin appendages.

**Fig. 4.**
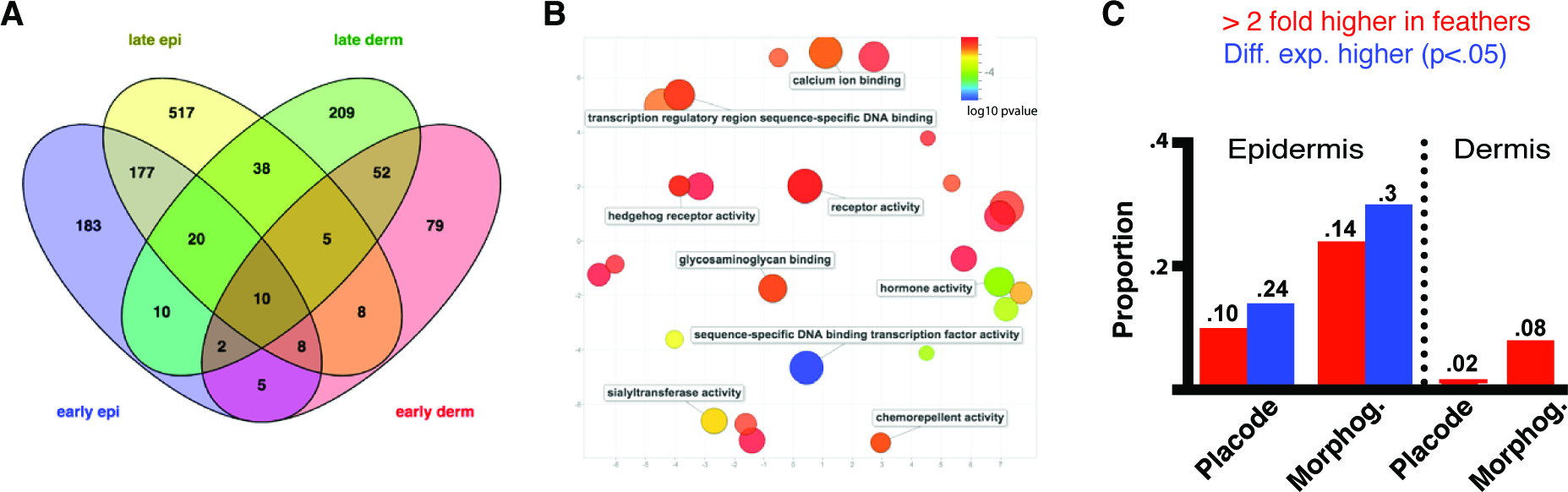
Candidate genes for feather developmental innovation. **(A)** Venn diagram showing extent to which feather candidate genes are shared between developmental stages and epidermis/dermis. **(B)** MDS plot of gene ontology terms showing morphogenesis stage feather candidate genes molecular function gene ontology terms. **(C)** Proportion of genes implicated previously in embryonic feather development by in-situ or immunostaining that are differentially expressed between chicken feathers and other chicken skin appendages.

We compared this new list of feather-specific genes to a list of 85 genes that have previously been shown to be expressed in embryonic feathers at similar developmental stages to those we sampled using in-situ hybridization or functional assays (table S3). Notably, we found little overlap between our candidate feather genes and those genes previously investigated in feather development (Fig. 4C). In particular, we found a number of known placode markers, such as *Bmp2* and *Fgf2* (7, 21, 22) that were not differentially expressed between feathers and scutate scales or other skin appendages. We also found a number of genes expressed in later stages of feather morphogenesis that were not specific to feathers in our avian dataset. This included the transcription factor *Dlx5* and adhesion molecule *NCAM1*, both of which exhibit localized expression patterns within developing feather buds (23). This result demonstrates that many previously described “feather genes”, particularly at the placode stage, actually play a more general role in avian skin appendage development.

### Development and evolution of feather-specific gene expression

We next sought to understand how feather-specific gene expression arises during development. For this, we quantified the change in expression of feather morphogenesis candidate genes during development, and compared it between feathers and other skin appendages. Surprisingly, we found that many of our feather morphogenesis candidates exhibited only a minor increase in expression on average during feather development (fig. 5A; mean ΔTPM = 1.7 +/-3.0 SEM). In contrast, average expression of these genes *decreased* during development of bird scutate scales (mean ΔTPM = ‐13.5 +/-1.7 SEM), and to a lesser extent in bird reticulate scales (mean ΔTPM = ‐8 +/-1.1 SEM). Alligator scales also showed decreased expression of these genes, although not to the same extent in as avian scutate scales, (symmetric scales mean ΔTPM = ‐2.1 +/-.73 SEM; asymmetric scales ΔTPM = ‐5.9 +/-1.8 SEM), suggesting that reduced expression of feather genes is a derived feature of avian scutate scales.

**Fig. 5.**
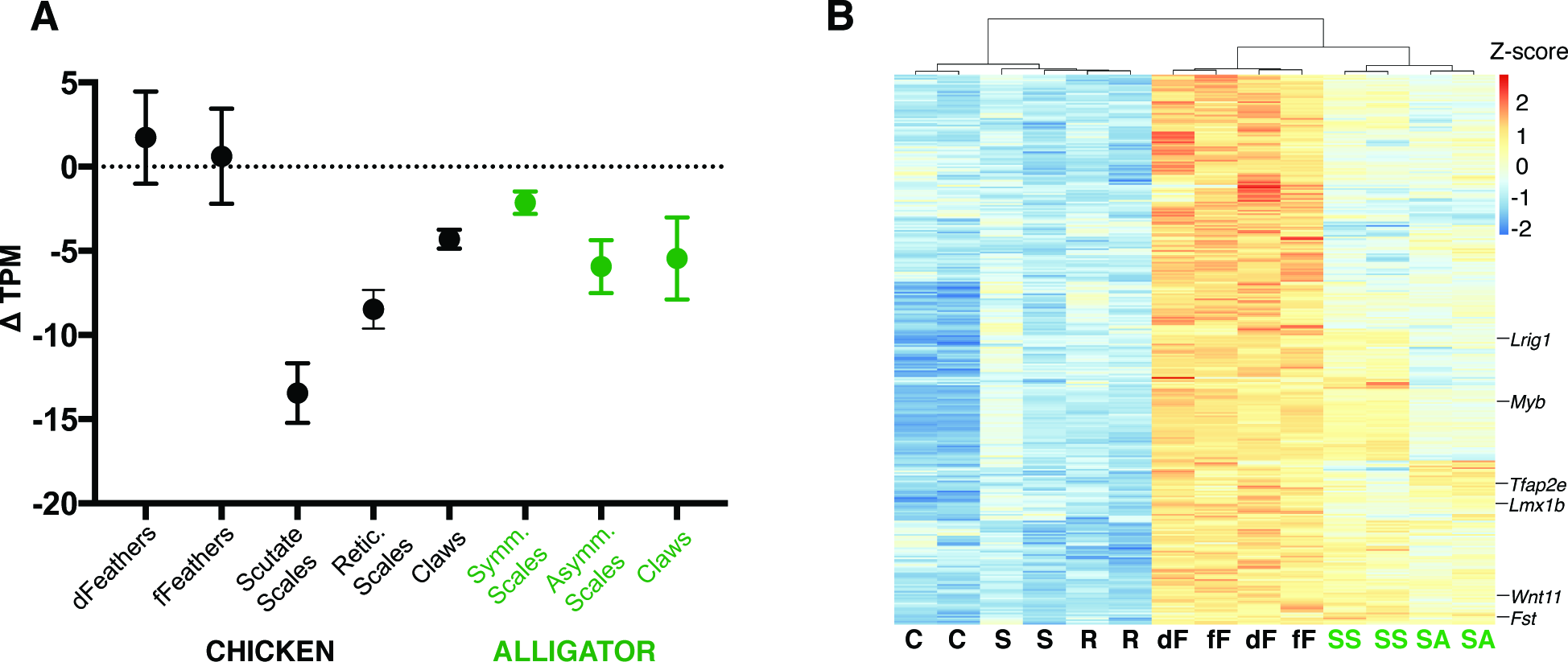
Development and evolution of feather candidate genes. **(A)** Mean change (+/- SEM) in relative expression of feather candidate genes from placode to morphogenesis stage. Scale bar is 1mm in A and B, 5mm in C. **(B)** Heatmap showing expression of morphogenesis stage feather candidate genes in other chicken and alligator skin appendages at the morphogenesis stage. C=chick claw, dF= chick dorsal feather, fF=chick femoral feather, R=chick reticulate scale, S=chick scutate scale, SA=alligator asymmetric scale, SS=alligator symmetric scale.

Notably, we also found that many of our feather morphogenesis candidate genes share more similar expression between feather and alligator scales than between bird and alligator scales at the morphogenesis stage (n = 142 of 329 one-to-one orthologs; fig. 5B and Supplementary Table 1). Among these were a variety of important feather transcription factors, such as *Myb*, which localizes to proliferation zones in developing feathers (24), as well as *Runx3* and *Mycn*, implicated in regulating flight feather development (14). Also included were genes from key signaling pathways implicated in orchestrating novel aspects of feather development, such as *Wnt11, Ptch1, Fst*, and *Fgf20* (7, 25-27). These findings suggest that genes important in feather morphogenesis were not necessarily recruited into the feather expression program *de novo*, but were in fact already expressed in ancestral archosaur scales. Their feather specific expression in bird feather morphogenesis was accomplished in part through the reduction of their expression in bird scutate scales. Thus, despite similar morphology, avian scutate scales have diverged transcriptionally from alligator scales, apparently in order to accommodate the reuse of ancestrally expressed genes in feathers. This is congruent with a previous study, which found that *Shh* and *Bmp2* are similarly expressed in feather, scutate scale, and alligator scale placodes, before being reutilized to organize barb development, a feather morphological novelty (7).

### The role of candidate genes in feather development

To explore the role of feather-specific genes in feather development, we characterized expression patterns for a subset of candidate genes (n=19) in chicken embryonic feathers (figs. 6A,B and figs. S3, S4). We selected candidate transcription factor and signaling pathway genes from both epidermis and dermis based on their likely role in regulating development, their differential expression between feathers and other avian skin appendages at the morphogenesis stage, and their conserved expression in both chicken and emu. We intentionally included two genes, *Ctbp2* and *Akap12*, that were specific to feathers only at the morphogenesis stage, but not at the placode stage (fig. 6C). We also investigated expression of two transcription factors (*Lmx1b* and *Tfap2e*), and a signaling pathway gene (*Lrig1*), that had similar expression between feathers and alligator scales.

**fig. 6.**
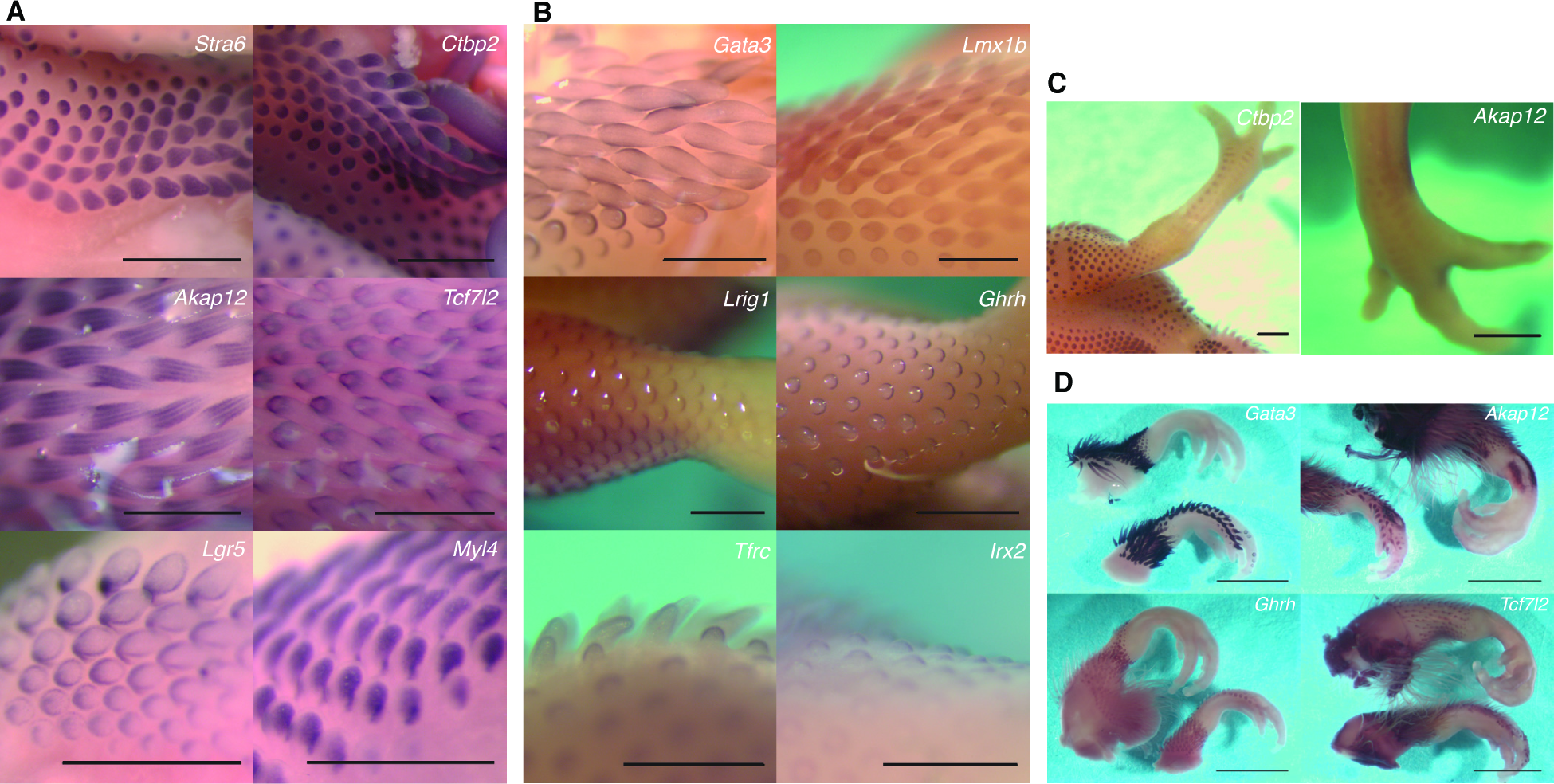
Feather Developmental Innovation. **(A and B)** Expression patterns in stage 36 chicken embryos of **(A)** dermal and **(B)** epidermal candidates for feather developmental innovation. **(C)** Morphogenesis stage feather candidates expressed in placode stage scutate scales. **(D)** Comparison of feather candidate gene expression in stage 38 Single Comb Leghorn chicken (top leg) and Silkie chicken (bottom leg).

Nearly all selected candidates (n=16 out of 19) exhibited localized expression to feathers, revealing three broad patterns for feather-specific genes. The first were genes expressed broadly across either feather bud epidermis (*Lmx1b* and *Tfap2e*) or dermis (*Ctbp2*). The two transcription factors, *Lmx1b* and *Tfap2e*, are also expressed in alligator scales. The second broad pattern were genes that exhibited localized expression within the developing feather bud. This was often associated with distinct feather developmental innovations, such as barb ridges (*Akap12*), the feather sheath (*Gata3*), and follicle (*Tcf7l2*). Notably, the dermal candidate gene *Akap12*, which is expressed in longitudinal stripes likely associated with developing barb ridges, is also expressed in the dermis of scutate scale placodes (fig 6C). This mirrors a similar pattern shown for the *Shh* and *Bmp2* in the epidermis, which were previously shown to be expressed in feather, scutate scale, and alligator scale placodes before organizing barb ridge development in later feather morphogenesis (7). Lastly, we found genes that showed polarized expression along both anterior-posterior (*Ghrh* and *Lrig1* in epidermis) and proximal-distal axes (*Irx2* and *Tfrc* in the epidermis and *Nr3c2, Stra6, Myl4* and *Lgr5* in the dermis). Several of these genes, such as *Ghrh, Lrig1*, and *Tfrc*, have been implicated in the control of cell proliferation (28), suggesting that the feather developmental program arose in part via the establishment of new developmental axes along which cell proliferation, and likely other developmental mechanisms, can be controlled by (29).

To further validate that our candidate feather genes were indeed specific to feathers, we investigated expression of four of these genes in the Silkie chicken breed (fig. 6D). Silkie chicken embryos simultaneously develop feathers and scutate scales on their feet, which enabled us to disentangle the effects of anatomical position and timing from true skin appendage identity. All four genes exhibited expression patterns in developing Silkie foot feather buds similar to embryonic feather buds in other regions, and were not expressed in neighboring scutate scales undergoing morphogenesis. These data confirm that these genes are highly specific to developing feathers, and their expression is not dependent upon the anatomical position of the skin appendage (fig. 6D).

## Discussion

### Model of skin appendage diversification

In this study, we investigated the evolution of feathers by comparing gene expression programs for skin appendages in three species of archosaurs: chick, emu and alligator. Based on our results, we propose an evolutionary model of skin appendage diversification that gave rise to the assemblage of modern archosaur skin organs (fig. 7). The similarity between feathers, avian scutate scales, and alligator body scales, relative to other skin appendages, suggests that these structures all derived from a body scale in the archosaur ancestor. This was likely similar to alligator body scales, with thickened placode epidermis and mature keratinized structure composed of β-keratin.

**fig. 7.**
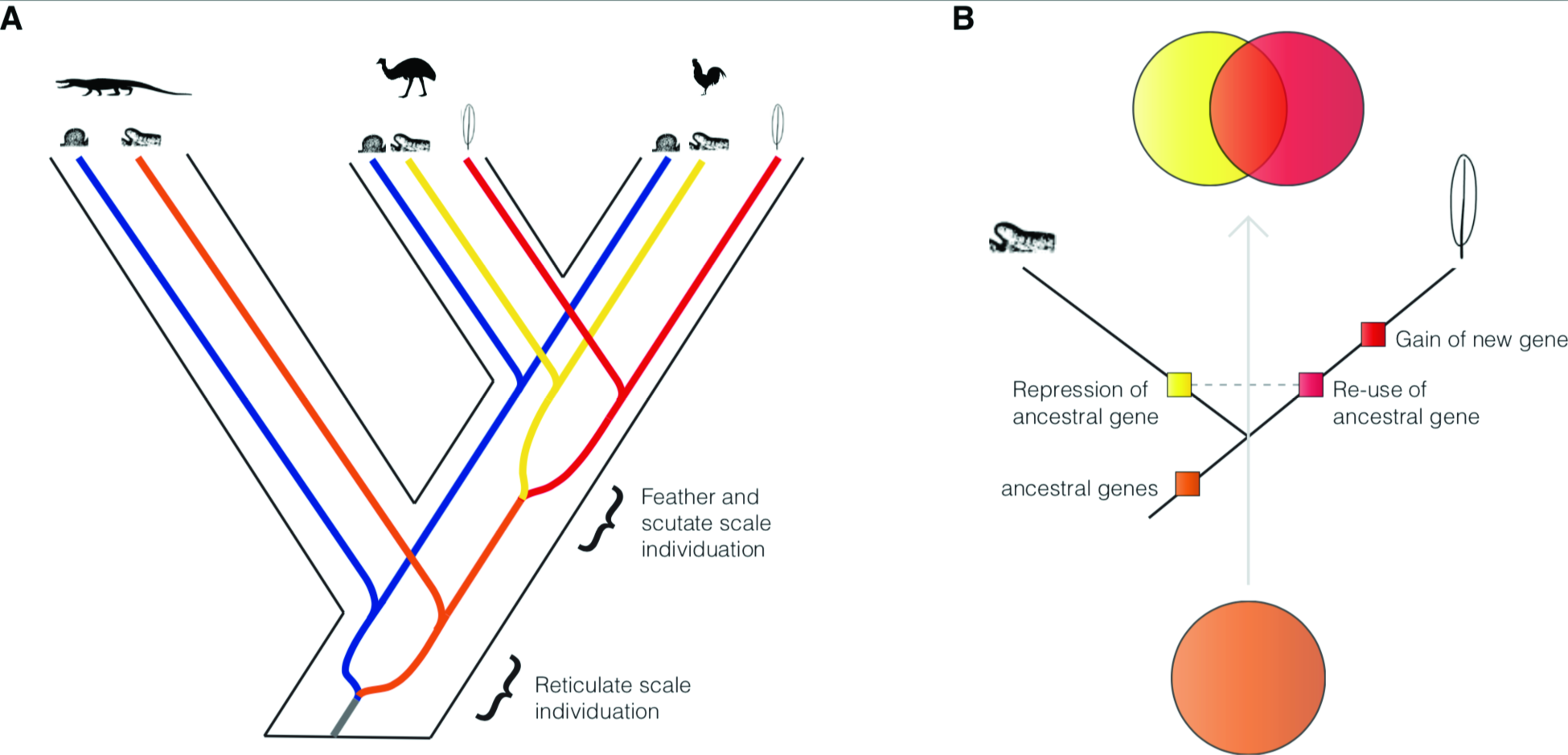
Evolutionary model of skin appendage diversification. **(A)** We propose that reticulate scales (blue lineage) individuated from other scale types prior to the archosaur ancestor. Feathers (yellow lineage) and avian scutate scales (orange lineage) then evolved in theropod dinosaurs from scales covering most of the ancestral archosaur (red lineage), which are still present broadly across the integument of American alligator. **(B)** Re-use and differential suppression of ancestr genes, and recruitment of new genes, are key processes during individuation of feathers and scutate scales.

Reticulate scales likely arose from a more ancient diversification event than scutate scales and feathers (chapter 9 in ref. 30). Although we did not sample reticulate scales in alligators, nearly all tetrapods, including alligators, exhibit distinct skin appendages on the ventral hindlimb footpads, and we suggest that the ancestral archosaur already had distinct reticulate scales on their foot pads and different body scales on the rest of the body. Moreover, reticulate scale identity depends on the transcription factor *engrailed*, a conserved marker of ventral limb identity across tetrapods (31-33). In birds, *engrailed* localizes to developing reticulate scales and represses *Shh* expression (32), a gene important in anterior-posterior polarization in feather, scutate scale, and alligator scale placodes (7). If reticulate scales originated from a more ancient evolutionary event, then we would predict that *engrailed* expression is localized to developing footpad scales of alligators and other non-avian reptiles.

In summary, our data support the following model (Fig. 7A): the ancestral archosaur already had reticulate scales on their foot pads and distinct body scales on the rest of the body. Within the theropod lineage, i.e. when feathers arose, the body scales differentiated into feathers and scutate scales, by subdividing the genes expressed in the ancestral body scale into two sets, some expressed in feathers and some expressed in scutate scales (Fig. 7B).

### Evolution of novelty by differential suppression of ancestrally-expressed genes

We found that the origination of feathers occurred in part through reuse of genes ancestrally-expressed in asymmetric body scales, accompanied by the downregulation of these genes in developing avian scutate scales. Thus, at the level of gene expression, both feathers and avian scutate scales represent derived skin appendages that arose via splitting an ancestral archosaur gene expression program. A similar process of evolutionary divergence has been described for the origination of novel cell types (34, 35), which occurred for instance when an ancestral photoreceptor diversified into distinct rods and cones, each specialized for different forms of light detection (34). This process has been referred to as *evolutionary individuation*, which emphasizes that the novel cell types must gain capacity to individually regulate and evolve their gene expression programs semi-independently of others. Importantly, our study shows that evolutionary individuation can also occur for more complex tissues and morphological structures. This result suggests that reuse and differential suppression of ancestrally expressed genes is a key mechanism by which new specialized tissues and structures can evolve.

In several respects, evolutionary individuation of cell types, tissues, and morphological structures is similar to diversification processes at the species and gene levels (36). In each case, distinct mechanisms result in the divergence of an ancestral evolutionary lineage – whether it be a gene, a cell type, an organ, a population, or a species – into multiple, independently evolving descendant lineages. For example, new species arise when an ancestral species is split into two, reproductively isolated sister species, via the formation of geographic barriers that isolate the two descendants, and through the subsequent evolution of reproductive incompatibilities. Likewise, gene duplication gives rise to two, independently evolving gene copies that are evolutionarily isolated due to independent inheritance occurring via semi-conservative replication during meiosis. In cell types and organs, divergence occurs through the evolution of regulatory mechanisms that enable the two descendant structures to maintain and evolve distinct gene expression patterns. Cell types and organs thus join other fundamental levels of biological organization capable of forming and diversifying as evolutionary lineages.

However, our study also indicates that evolutionary individuation of tissues and morphological structures does not immediately result in their complete evolutionary independence. For instance, we found that feathers and scutate scales exhibit highly correlated gene expression evolution at both placode and morphogenesis stages. This suggests that two related tissues can exhibit distinct and highly specialized morphology, yet still retain relatively limited ability to differentially evolve their gene expression programs. This stands in contrast to estimates of correlated evolution made for other organs, such as liver in mammals, which was found to evolve gene expression nearly independently of other mammalian tissues (20). Thus, although individuation of organismal parts such as cell types or organs may result in their complete evolutionary independence, this is not always necessary for the formation of morphologically distinct structures, as illustrated by our findings for feathers and avian scutate scales. Incomplete evolutionary independence has also been noted at other levels of biological organization, such as when genes undergo concerted evolution due to gene conversion (37).

## Conclusion

The origin of feathered theropod dinosaurs represents a dramatic transition in animal evolution. Over the last two decades, discoveries of feathered non-avian dinosaurs has fueled a paleontological revolution, uncovering key evolutionary steps on the path to modern birds (1). For instance, fossil discoveries from the Jurassic have shown that the first feathers were likely simple filamentous structures, only later acquiring the complex branched structure necessary for supporting flight (1).

We have used comparative transcriptomics to investigate the evolutionary emergence of feathers and bird scales (36, 38). We show that feathers and scutate scales evolved when ancestral archosaur scales on different parts of the body became distinct, individualized structures. Our results show that the individuation process giving rise to feathers and bird scutate scales occurred when ancestral scale genes were reprogrammed for feather development, and repressed in avian scutate scales. This process of individuation is analogous to other diversification processes in evolution, such as speciation and gene duplication, which give rise to new species and genes. In each case, an ancestral entity evolves to become two distinct and independently, or partially-independently, evolving entities.

## Data availability

RNAseq data that support the findings of this study may be obtained by request from the corresponding author.

## Code availability

All code used to generate results in this paper is available from the authors upon request.

## Acknowledgements

We thank Peter Brazaitis, Walter Brenckle, Ruth Elsey and the staff of Louisiana Rockefeller Wildlife Refuge, Brendan Murphy, Philip Musser, John VandenBrooks, Zhe Wang, Myrna Watanabe, Gregory Watkins-Colwell, and Rebecca Young for helping with various aspects of this project. This research was supported by the National Science Foundation Graduate Research Fellowship Program under Grant No. DGE-1122492, the W.R. Coe Funds from Yale University, and the Dillon and Mary Ripley Graduate Fellowship Fund at Yale University (JMM).

---

**Fig. S1.**
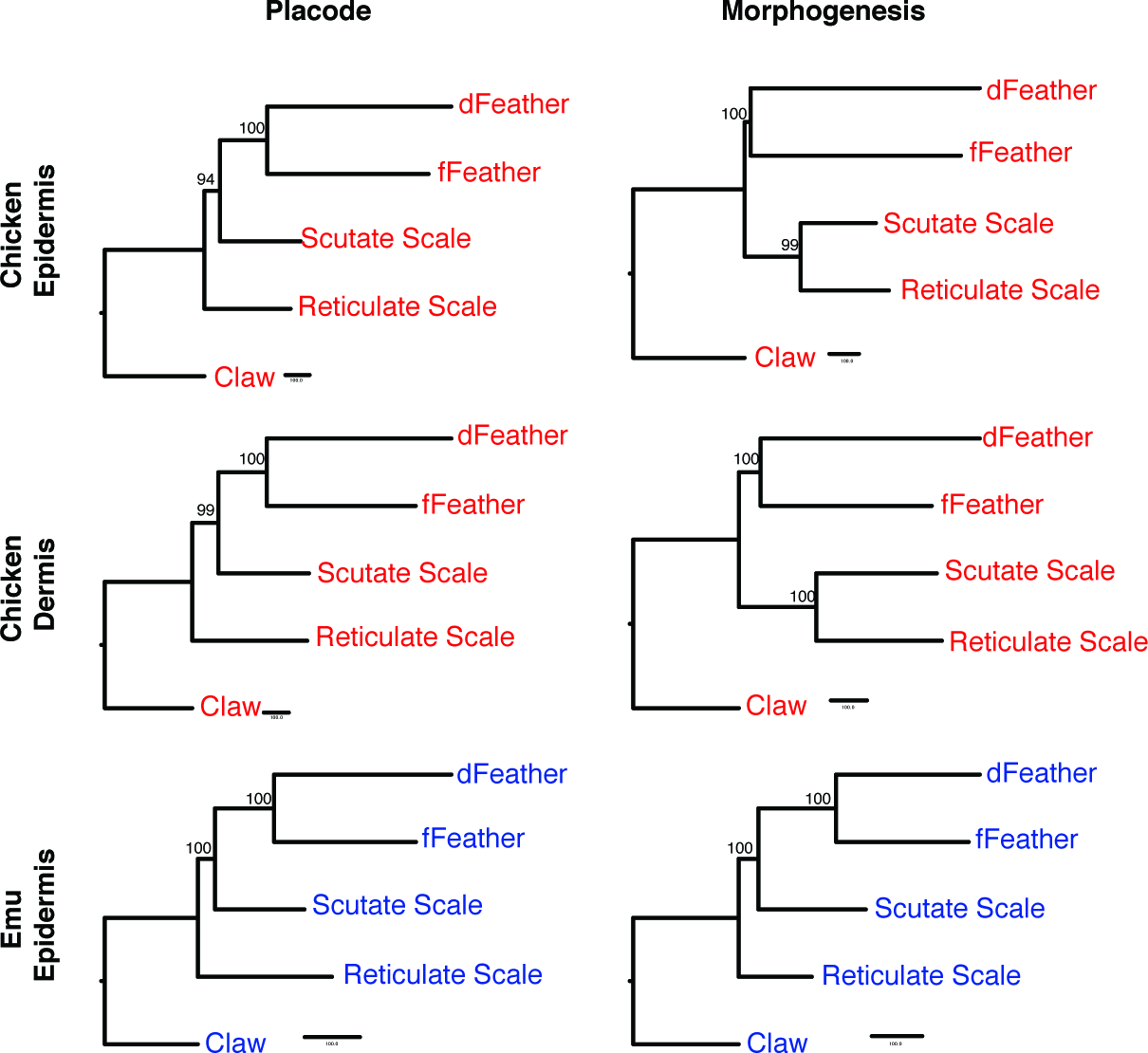
Parsimony tree reconstructions of chicken (red) and emu (blue) skin appendage relationships. Trees were constructed using discretized expression values. Support values were calculated with 10,000 bootstrap replicates.

**Fig. S2.**
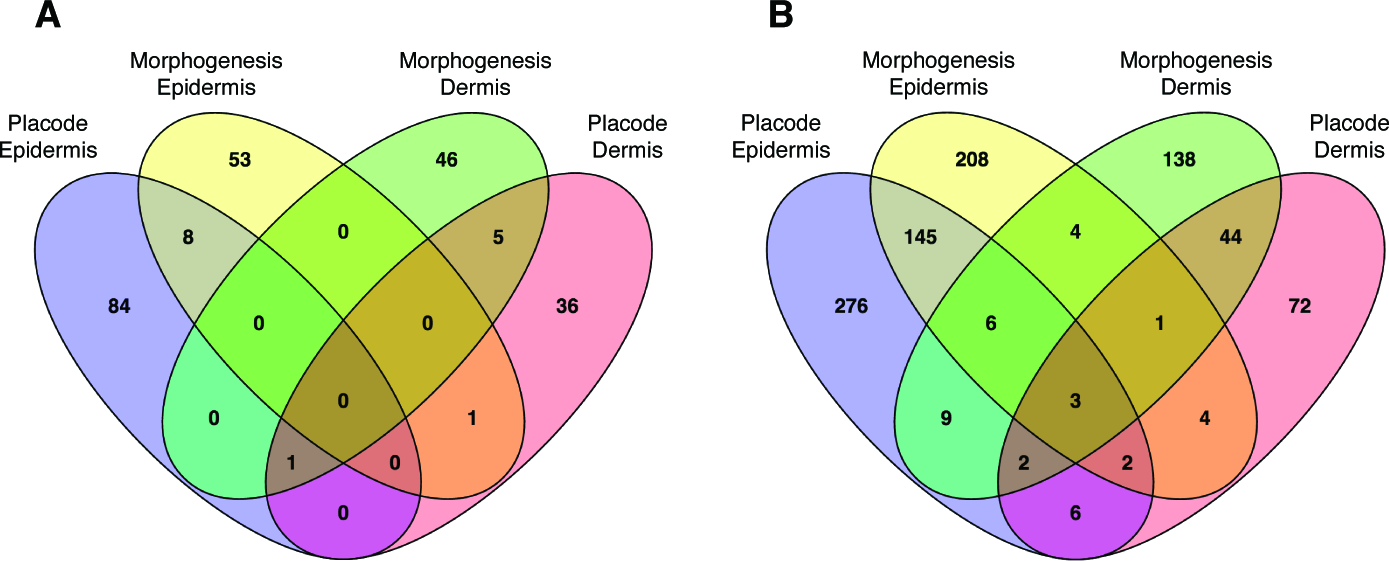
Chicken skin appendage synapomorphies. (A) Expressed genes shared between feathers and scutate scales. (B) Expressed genes shared between feathers, scutate scales, and reticulate scales.

**Fig. S3.**
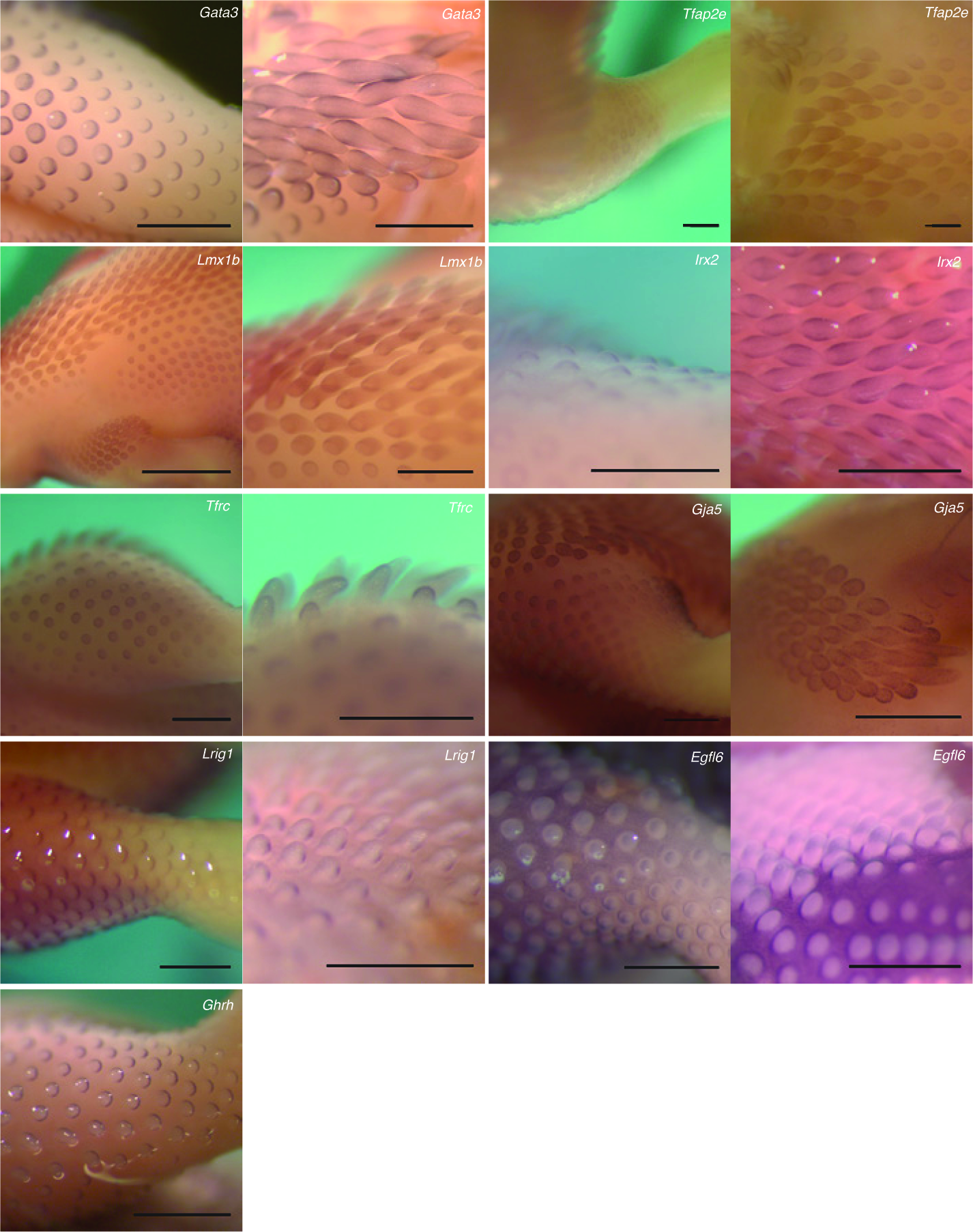
In situ hybridization of epidermal candidate feather genes in whole mount chick. Expression patterns in day 10 embryonic chicken are shown for two feather tracts at either the placode/early bud stage (left panel) or during feather bud morphogenesis (right panel). *Ghrh* expression (single panel) is visible only in early and mid stage feather buds. Scale bar is 1mm in all panels except 5mm in left *Lmx1b* panel.

**Fig. S4.**
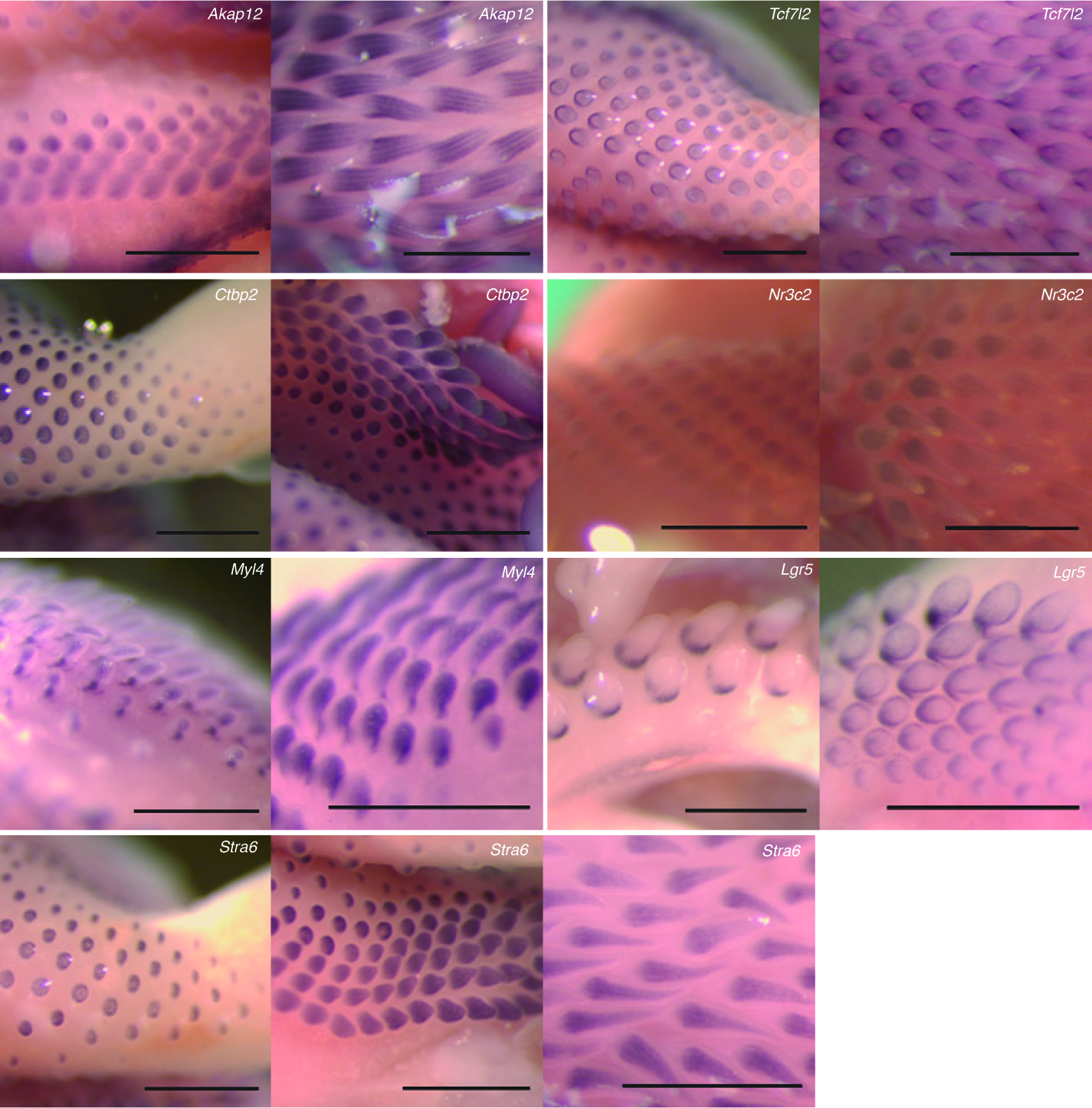
In situ hybridization of dermal candidate feather genes in whole mount chick. Expression patterns in day 10 embryonic chicken are shown for two feather tracts at either the placode/early bud stage (left panel) or during feather bud morphogenesis (right panel). *Stra6* expression is shown in three panels, illustrating progressive stages of feather development from left to right. Scale bar is 1mm in all panels.

**Table S1.** Estimates and 95% confidence intervals of correlated evolution (*γ*) between epidermal transcriptomes of avian skin appendages. p=placode stage, m=morphogenesis stage.

|  | Scutate Scales |  | Reticulate Scales |  | Claws |  |
| --- | --- | --- | --- | --- | --- | --- |
|  | p | m | p | m | p | m |
| *Feathers* | 0.88 [0.87, 0.89] | 0.84 [0.83, 0.85] | 0.79 [0.78, 0.81] | 0.81 [0.80, 0.82] | 0.71 [0.70, 0.73] | 0.70 [0.68, 0.71] |
| *Scutate Scales* |  |  | 0.86 [0.85, 0.87] | 0.87 [0.86, 0.88] | 0.72 [0.70, 0.74] | 0.70 [0.68, 0.72] |
| *Reticulate Scales* |  |  |  |  | 0.71 [0.69, 0.72] | 0.74 [0.72, 0.75] |

**Table S2.** Gene ontology enrichment of feather candidate genes at the morphogenesis stage. Shown are all “molecular function” gene ontology terms with p-values <.01 (corrected for multiple testing).

| Gene Ontology ID | Gene Ontology Category | P-value | # Diff. Exp. | # In Category |
| --- | --- | --- | --- | --- |
| GO:0043565 | sequence-specific DNA binding | 1.3E-07 | 58 | 436 |
| GO:0003700 | sequence-specific DNA binding transcription factor activity | 2.4E-06 | 70 | 597 |
| GO:0005179 | hormone activity | 1.1E-04 | 12 | 58 |
| GO:0008179 | adenylate cyclase binding | 1.3E-04 | 4 | 5 |
| GO:0005158 | insulin receptor binding | 1.5E-04 | 7 | 19 |
| GO:0004800 | thyroxine 5'-deiodinase activity | 6.1E-04 | 3 | 4 |
| GO:0008373 | sialyltransferase activity | 9.3E-04 | 7 | 25 |
| GO:0005159 | insulin-like growth factor receptor binding | 1.4E-03 | 4 | 9 |
| GO:0003677 | DNA binding | 2.3E-03 | 91 | 1012 |
| GO:0005509 | calcium ion binding | 2.3E-03 | 52 | 499 |
| GO:0001077 | RNA polymerase II core promoter proximal region sequence-specific DNA bindi | 2.8E-03 | 8 | 37 |
| GO:0045499 | chemorepellent activity | 2.9E-03 | 3 | 5 |
| GO:0035240 | dopamine binding | 3.1E-03 | 3 | 5 |
| GO:0005539 | glycosaminoglycan binding | 3.3E-03 | 5 | 15 |
| GO:0001965 | G-protein alpha-subunit binding | 3.4E-03 | 3 | 5 |
| GO:0003705 | RNA polymerase II distal enhancer sequence-specific DNA binding... | 3.7E-03 | 8 | 38 |
| GO:0000976 | transcription regulatory region sequence-specific DNA binding | 3.9E-03 | 7 | 33 |
| GO:0004364 | glutathione transferase activity | 3.9E-03 | 4 | 11 |
| GO:0005102 | receptor binding | 4.3E-03 | 22 | 175 |
| GO:0008158 | hedgehog receptor activity | 5.0E-03 | 4 | 11 |
| GO:0048256 | flap endonuclease activity | 5.4E-03 | 2 | 2 |
| GO:0097108 | hedgehog family protein binding | 5.4E-03 | 2 | 2 |
| GO:0005543 | phospholipid binding | 6.4E-03 | 28 | 236 |
| GO:0004872 | receptor activity | 6.6E-03 | 15 | 103 |
| GO:0046982 | protein heterodimerization activity | 7.8E-03 | 23 | 206 |
| GO:0019955 | cytokine binding | 8.9E-03 | 3 | 7 |
| GO:0008146 | sulfotransferase activity | 8.9E-03 | 8 | 45 |
| GO:0004198 | calcium-dependent cysteine-type endopeptidase activity | 8.9E-03 | 4 | 12 |
| GO:0005057 | receptor signaling protein activity | 9.3E-03 | 4 | 13 |

## Additional data table S3 (separate file)

Detailed information on tissue samples, previously described feather genes, and primers for in situ probes.

### Detailed Materials and Methods

#### Collection of tissue samples

We investigated gene expression in skin appendages of three diverse archosaur species: chicken (*Gallus gallus*), emu (*Dromaius novaehollandiae*), and American alligator (*Alligator mississipiensis*) (Fig. 1a). Chicken and emu are members of the neognaths and paleognaths, respectively, the two oldest clades in living birds. American alligator is a crocodilian, the reptile clade most closely related to living birds (Fig. 1a).

Fertile eggs of Single Comb White Leghorn chicken were obtained from the University of Connecticut Poultry Farm. We obtained Silkie chicken eggs from an independent breeder in New York. Emu eggs were obtained from the Songline Emu Farm in Gill, MA. Bird eggs were incubated with rotation in a G.Q.F 1502 Sportsmen Incubator at 37.8°C (chicken) or 35.5°C (emu). Alligator eggs were collected from the Rockefeller State Wildlife Refuge in Grand Chenier, LA under the direction of refuge manager Ruth Elsey. A detailed description of alligator egg incubation is provided elsewhere (8, 39).

We dissected developing skin in chicken and emu embryos from five different locations in chicken and emu: dorsal, femoral (chicken only), and flank (emu only) feather tracts, scutate scales from the dorsal tarsometatarsus adjacent to the base of the digits, reticulate scales on the ventral metatarsal footpad, and claws from the hindlimb digit tips (Fig. 1b). We dissected developing skin in alligator from three locations: asymmetric scales on the dorsal thigh, symmetric scales on the flank, and claws from the tips of the digits (Fig. 1b). For each dissection we enzymatically separated epidermis and dermis by placing the tissue in a minimal amount of 10 mg/mL dispase at 4°C for approximately 13 hours and then separating with forceps. We confirmed this approach yielded clean separation by embedding separated skin in OCT compound and sectioning on a cryostat. We also verified that tissues did not express known markers of the apposing tissue. Both approaches confirmed the absence of cross-contamination.

For each skin appendage we sampled two developmental stages: a “placode” stage and a later “morphogenesis” stage. Skin appendages develop heterogeneously across the organism (40, 41), and take different amounts of time to undergo morphogenesis (3, 4, 42). For the placode stage we sampled tissue at the point when the skin appendage first became visible on the skin surface (Fig. 1b and Supplementary Table 1). For the morphogenesis stage, we sampled tissue at the point when each tissue began to undergo its unique morphogenesis, as assessed by sectioning and staining with hematoxylin and eosin (Fig. 1b and Supplementary Table 1). To verify that we had collected similar stages of development for each replicate and across species we collected tissue for RNAseq from the left side of the embryo, and sectioned and visualized tissue from the embryo’s right side.

#### mRNA sequencing and normalization

We extracted RNA from individual skin samples using a Qiagen Rneasy kit and verified quality of the RNA on an Agilent 2100 Bioanalyzer (Supplementary Table 1). Strand-specific polyadenylated RNA libraries were prepared for sequencing by the Yale Center for Genome Analysis using an in-house protocol. We pooled 4 samples per Illumina Hiseq 2000 flow cell lane, yielding 30-50 million reads per sample. For each epidermal sample we sequenced two biological replicates, except for placode stage emu reticulate scales and claw, for which we obtained only one replicate due to limitations in the number of emu embryos we could obtain at the appropriate stage. For dermis, we sequenced one replicate for each skin appendage in chicken.

We first assessed quality of the sequenced reads using the program fastQC (http://www.bioinformatics.babraham.ac.uk/projects/fastqc/). Reads were then mapped to their respective species genome currently available at the time: WASHUC2 for chicken (Ensembl release 68 downloaded October 4^th^, 2012), allmis1 for alligator (downloaded from the Crocodile Genome Project on December 4^th^, 2012), and a preliminary version of the Emu genome generated by the Alan Baker Lab at the University of Toronto (mapped by Alison Cloutier on December 5^th^, 2012). We mapped reads using Tophat2 v2.0.6 using the –GTF option, and assigned mapped reads to genes with HTSeq v0.5.3p9(43) implemented in python v2.7.2. For HTSeq we set the option to require that a read maps to the correct strand, and used the “intersection-nonempty” option for handling reads mapped to multiple genes. We then estimated transcripts per million (TPM) relative expression values independently for each species, and excluded genes on the chicken W chromosome to remove any signal based on the sex of the sample.

Comparisons across species are challenging because relative expression values may be biased in a species-specific manner due to differences in gene number, annotation quality, and correlated changes in gene expression that occur across all skin tissue (20, 36). In our initial clustering attempts, we found that these biases resulted in a strong tendency for samples to cluster by species, even for tissues with unambiguous homology, such as claw. This was not related to sequencing batch, as samples sequenced on the same lane, but from different species, still clustered strongly by species. To overcome these biases, and extract information about tissue homology, we implemented a three-step scaling and normalization procedure that enabled comparing transcriptomes across species, and recovered claw as a homologous group. First, we applied scaling factors to emu (.55x) and alligator (.62x) expression values calculated using the method of Musser and Wagner (36). This corrects for differences in relative expression values between species due solely to differences in the number of annotated genes (which affects the denominator of the TPM calculation). Second, we applied a square root transformation to remove the dependence of the mean on variance. Finally, we used the ComBat function(44) in the R library “sva” to remove species-specific effects caused by differences in genome quality, read mappability, and correlated evolutionary changes across all skin samples. This function was applied separately to scaled TPM expression matrices containing either early or late epidermal skin samples prior to conducting the following cross-species analyses: hierarchical clustering, t-SNE, and heatmaps. In each case, we removed genes with zero variance and designated the species a tissue came from as its “batch”. We did not specify covariates (e.g. tissue type) to avoid making assumptions about tissue homology.

#### mRNA-seq analyses

For analyses in which we compared expression values across species we used ComBat corrected datasets. We conducted hierarchical clustering analysis independently for early and late epidermal samples from all three species using the R package “pvclust” with default values and 10,000 boostrap replicates. For both early and late epidermal datasets, we conducted T-distributed stochastic neighbor embedding analyses using the “Rtsne” package in R.

Phylogenetic maximum parsimony trees were built independently for each species using the phylogenetic software paup* v4.0b10 with the alltrees. This analysis required generating a discretized dataset, in which genes were called “on” or “off’. Because ComBat alters each gene independently, it is impossible to apply a uniform cutoff for our ComBat corrected dataset. Hence, we limited our parsimony analyses to TPM matrices from each species independently, without applying the ComBat correction. For this we used a cutoff of 3 TPM for a gene to be called “on” by using the method of Wagner et al.(45), which estimates a cutoff that corresponds to qualitative differences in chromatin conformation(36, 46, 47). To reduce the impact of expression noise around the cutoff we required that both replicates express a gene above 3 TPM for that gene to be called “on” in that tissue. We assigned claw as the outgroup and justified this choice based on *a priori* evidence that claws do not express many genes known to be shared by feathers and scales, and because hierarchical clustering confirmed claw transcriptomes were highly divergent from feathers and scales in both chicken and emu. Node support was assessed by generating 10000 bootstrap replicates.

We assessed correlated gene expression evolution among pairs of bird skin appendages using the method of Liang et al. (20), which estimates gamma as the pearson correlation between gene expression contrast vectors for two tissues in two species. For these estimates, we used only one-to-one orthologs shared between chicken and emu, and used TPM values that had been scaled and square-root transformed, but not normalized with ComBat, so as not to remove any correlated evolution signal. These comparisons were limited to epidermal tissue as they required having samples from both chicken and Emu. For each skin appendage, we averaged expression across replicates, and then calculated the contrast vector by taking the difference in expression between chicken and emu. Following Liang et al. (20), we estimated correlated evolution as the pearson correlation between pairs of skin appendage contrast vectors, and conducted bootstrapping (n=1000 bootstrap replicates) to estimate the 95% confidence interval.

To identify new candidate genes for feather developmental innovation we used the criteria that a gene must be expressed greater than 2-fold higher in feathers compared to other bird skin appendages. In epidermis, these genes represented a subset of the genes that were differentially expressed between feathers and other skin appendages, and were thus a more conservative set of candidate genes than those differentially expressed. Furthermore, using fold difference as our criteria, rather than statistical significance, allowed us to compare epidermal and dermal candidate genes, since we only had 1 replicate for dermal tissue samples. For morphogenesis stage candidate genes we conducted gene ontology analysis using the R bioconductor 3.1 package goseq(48) and visualized using the online software REVIGO(49) implemented online (http://revigo.irb.hr/), which clusters related gene ontology terms.

We also conducted a thorough review of the literature to identify genes previously described as being expressed in feather epidermis or dermis at the stages for which we assayed gene expression (Supplementary Table 1). For these, we required expression to have been documented using either in-situ hybridization or immunohistochemistry in chicken embryonic feathers. Furthermore, the gene must exhibit localized expression within the feather bud epidermis and/or dermis. In a few cases we included genes that were localized to neighboring interfeather epidermis/dermis. We compared expression of these genes in our dataset using differential expression analysis as implemented in the R package DESeq2(50), using raw read counts as required by this method for differential expression. Differentially expressed genes were identified between chicken feathers and each other chicken skin appendage independently. For a gene to be considered “feather-specific” we required that it was differentially expressed higher versus each other chicken skin appendage. For those genes that were feather-specific in chicken at the morphogenesis stage, and had one-to-one orthologs in alligator, we visualized and compared their expression in the two species using the “pheatmap” function in R. For this we applied the scale function in R to our ComBat normalized dataset to yield z-scores for all feather-specific (in chicken) one-to-one orthologs (between chicken and alligator).

#### In situ hybridization

We explored spatial patterns of expression for 19 candidate feather genes by conducting whole mount in-situ hybridization (WMISH) on stage 36(41) chick embryos, which allowed us to observe expression patterns in a variety of different feather development stages, as well as in early scutate and claw development. We developed riboprobes for WMISH using a pcr-based approach (Supplementary Table 1). For this we amplified the target region from chicken cDNA using gene-specific primers (Supplementary Table 1), followed by an additional round of amplification using a forward (for sense) or reverse (for antisense) primer composed of a T3 RNA polymerase promoter sequence 5’ of the gene-specific primer sequence. We verified successful amplification of the target region by gel electrophoresis and sanger sequencing. The PCR product was then purified and used to generate DIG-labeled probes in a transcription reaction with either T3 or T7 RNA polymerases. Our WMISH protocol followed that of GEISA(51) (Gallus Expression in Situ Hybridization Analysis). We performed a minimum of 2 replicate WMISH experiments in stage 36 embryos for each antisense probe and 1 replicate for each sense probe.

## References

1. Xu X, et al. (2014) An integrative approach to understanding bird origins. Science 346(6215): 1253293.

2. Chen C-F, et al. (2015) Development, Regeneration, and Evolution of Feathers. Annu Rev Anim Biosci 3(1): 169–195.

3. Sawyer RH (1972) Avian scale development. I. Histogenesis and morphogenesis of the epidermis and dermis during formation of the scale ridge. J Exp Zool 181(3):365–383.

4. Sawyer RH, Craig KF (1977) Avian scale development. Absence of an “epidermal placode” in reticulate scale morphogenesis. J Morphol 154(1):83–93.

5. Prum R (1999) Development and evolutionary origin of feathers. J Exp Zool 285(4):291–306.

6. Lucas AM, Stettenheim PR (1972) Avian Anatomy Integuments Part I, II (US Gov. Print. Off., Washington, DC).

7. Harris M, Fallon J, Prum R (2002) Shh-Bmp2 signaling module and the evolutionary origin and diversification of feathers. J Exp Zool 294(2):160–176.

8. Musser JM, Wagner GP, Prum RO (2015) Nuclear β-catenin localization supports homology of feathers, avian scutate scales, and alligator scales in early development. Evol Dev 17(3):185–194.

9. Di-Poi N, Milinkovitch MC (2016) The anatomical placode in reptile scale morphogenesis indicates shared ancestry among skin appendages in amniotes. Science Advances 2(6):e1600708–e1600708.

10. Dhouailly D, Hardy MH, Sengel P (1980) Formation of feathers on chick foot scales: a stage-dependent morphogenetic response to retinoic acid. J Embryol Exp Morphol 58:63–78.

11. Yu M, Wu P, Widelitz R, Chuong C (2002) The morphogenesis of feathers. Nature.

12. Wu P, et al. (2017) Multiple Regulatory Modules Are Required for Scale-to-Feather Conversion. Mol Biol Evol 35(2):417–430.

13. Li A, et al. (2017) Diverse feather shape evolution enabled by coupling anisotropic signalling modules with self-organizing branching programme. Nat Commun 8:1–13.

14. Ng CS, et al. (2015) Transcriptomic analyses of regenerating adult feathers in chicken. BMC Genomics 16(1):756.

15. Chang K-W, et al. (2015) Emergence of differentially regulated pathways associated with the development of regional specificity in chicken skin. BMC Genomics 16(1):22.

16. Bao W, Greenwold MJ, Sawyer RH (2016) Expressed miRNAs target feather related mRNAs involved in cell signaling, cell adhesion and structure during chicken epidermal development. Gene 591(2):393–402.

17. Li A, et al. (2013) Shaping organs by a wingless-int/Notch/nonmuscle myosin module which orients feather bud elongation. P Natl Acad Sci Usa 110(16):E1452–61.

18. Yue Z, Jiang TX, Wu P, Widelitz RB, Chuong C-M (2012) Sprouty/FGF signaling regulates the proximal–distal feather morphology and the size of dermal papillae. Dev Biol 372(1):45–54.

19. Lynch VJ, et al. (2008) Adaptive changes in the transcription factor HoxA-11 are essential for the evolution of pregnancy in mammals. Proc Natl Acad Sci USA 105(39): 14928–14933.

20. Liang C, Musser JM, Cloutier A, Prum RO, Wagner GP (2018) Pervasive Correlated Evolution in Gene Expression Shapes Cell and Tissue Type Transcriptomes. Genome Biol Evol 10(2):538–552.

21. Song H, Lee S, Goetinck P (2004) FGF-2 signaling is sufficient to induce dermal condensations during feather development. Dev Dynam 231(4):741–749.

22. Scaal M, et al. (2002) BMPs induce dermal markers and ectopic feather tracts. Mech Develop 110(1-2):51–60.

23. Rouzankina I, Abate-Shen C, Niswander L (2004) Dlx genes integrate positive and negative signals during feather bud development. Dev Biol 265(1):219–233.

24. Desbiens X, Queva C, Jaffredo T, Stehelin D, Vandenbunder B (1991) The relationship between cell proliferation and the transcription of the nuclear oncogenes c-myc, c-myb and c-ets-1 during feather morphogenesis in the chick embryo. Development 111(3):699.

25. Chodankar R, et al. (2003) Shift of localized growth zones contributes to skin appendage morphogenesis: role of the Wnt/beta-catenin pathway. J Invest Dermatol 120(1):20–26.

26. Abate-Chang C-H, et al. (2004) Distinct Wnt members regulate the hierarchical morphogenesis of skin regions (spinal tract) and individual feathers. Mech Develop 121(2): 157–171.

27. Ohyama A, Saito F, Ohuchi H, Noji S (2001) Differential expression of two BMP antagonists, gremlin and Follistatin, during development of the chick feather bud. Mech Develop 100(2):331–333.

28. O’Donnell KA, et al. (2006) Activation of Transferrin Receptor 1 by c-Myc Enhances Cellular Proliferation and Tumorigenesis. Molecular and Cellular Biology 26(6):2373–2386.

29. Prum R (2005) Evolution of the morphological innovations of feathers. J Exp Zool 304(6):570–579.

30. Wagner GP (2014) Homology, genes, and evolutionary innovation (Princeton University Press, Princeton, NJ).

31. Loomis CA, et al. (1996) The mouse Engrailed-1 gene and ventral limb patterning. Nature 382(6589):360–363.

32. Prin F, Logan C, D'Souza D, Ensini M, Dhouailly D (2004) Dorsal versus ventral scales and the dorsoventral patterning of chick foot epidermis. Dev Dynam 229(3):564–578.

33. Christen B, Slack JM (1998) All limbs are not the same. Nature 395(6699):230–231.

34. Arendt D (2008) The evolution of cell types in animals: emerging principles from molecular studies. Nat Rev Genet 9(11):868–882.

35. Arendt D, et al. (2016) The origin and evolution of cell types. Nat Rev Genet 17(12):744–757.

36. Musser JM, Wagner GP (2015) Character trees from transcriptome data: Origin and individuation of morphological characters and the so-called “species signal.” J Exp Zool (Mol Dev Evol) 324(7):588–604.

37. Coen E, Strachan T, Dover G (1982) Dynamics of concerted evolution of ribosomal DNA and histone gene families in the melanogaster species subgroup of Drosophila. Journal of Molecular Biology 158(1): 17–35.

38. Kin K (2015) Inferring cell type innovations by phylogenetic methods-concepts, methods, and limitations. J Exp Zool B Mol Dev Evol 324(8):653–661.

39. Vargas AO, Kohlsdorf T, Fallon JF, VandenBrooks J, Wagner GP (2008) The Evolution of HoxD-11 Expression in the Bird Wing: Insights from Alligator mississippiensis. PLoS One 3(10):e3325.

40. Mayerson PL, Fallon JF (1985) The spatial pattern and temporal sequence in which feather germs arise in the white Leghorn chick embryo. Dev Biol 109(2):259–267.

41. Hamburger V, Hamilton HL (1951) A series of normal stages in the development of the chick embryo. J Morphol 88(1):49–92.

42. Wessells NK (1965) Morphology and proliferation during early feather development. Dev Biol 12(1): 131–153.

43. Anders S, Pyl PT, Huber W (2015) HTSeq‐‐a Python framework to work with high-throughput sequencing data. Bioinformatics 31(2): 166–169.

44. Johnson WE, Li C, Rabinovic A (2007) Adjusting batch effects in microarray expression data using empirical Bayes methods. Biostatistics 8(1): 118–127.

45. Wagner GP, Kin K, Lynch VJ (2013) A model based criterion for gene expression calls using RNA-seq data. Theor Biosci 132(3): 159–164.

46. Hebenstreit D, et al. (2011) RNA sequencing reveals two major classes of gene expression levels in metazoan cells. Molecular Systems Biology 7:497.

47. Kin K, Nnamani MC, Lynch VJ, Michaelides E, Wagner GP (2015) Cell-type Phylogenetics and the Origin of Endometrial Stromal Cells. Cell Reports 10(8): 1398–1409.

48. Young MD, Wakefield MJ, Smyth GK, Oshlack A (2010) Gene ontology analysis for RNA-seq: accounting for selection bias. Genome Biology 11(2):R14.

49. Supek F, Bošnjak M, Škunca N, Šmuc T (2011) REVIGO Summarizes and Visualizes Long Lists of Gene Ontology Terms. PLoS One 6(7):e21800.

50. Anders S, Huber W (2010) Differential expression analysis for sequence count data. Genome Biology 11(10):R106.

51. Darnell DK, et al. (2007) GEISHA: an in situ hybridization gene expression resource for the chicken embryo. Cytogenet Genome Res 117(1-4):30–35.

